# Modulation of Common Synaptic Inputs and Motor Unit Recruitment Threshold in Triceps Surae Muscles: Effects of Ankle Position

**DOI:** 10.1101/2024.12.16.628556

**Authors:** Xin Sienna Yu, Jackson T. Levine, José L. Pons

**Affiliations:** Legs and Walking Lab, Shirley Ryan AbilityLab, Chicago, IL, 60611, USA; Department of Physical Medicine and Rehabilitation, Northwestern University, Chicago, IL, USA; Department of Biomedical Engineering, Northwestern University, Evanston, IL, USA

**Author notes:** These authors have contributed equally to the work (Xin Sienna Yu, Jackson T. Levine). **Corresponding author address:** Xin Sienna Yu, PhD, Legs + Walking Lab, Shirley Ryan AbilityLab, Floor 23, 355 E Erie St, Chicago, IL 60611, USA. **email address:**.

**Keywords:** Motor unit, muscle length, common synaptic input, force fluctuations, recruitment threshold

## Abstract

The objective of this study was to investigate how altering muscle length by changing ankle position affects force control, motor unit recruitment, and motor unit coherence within and across triceps surae muscles. Sixteen healthy young adults performed isometric plantarflexion (PF) at three ankle positions with the ankle plantarflexed at 20° (PF20°), at the neural position (PF0°) and dorsiflexed at 20° (DF20°). High-density surface electromyography was used to record the medial and lateral heads of the gastrocnemius muscle (GM and GL), and the medial and lateral portions of the soleus muscle (SL and SM). Motor unit cumulative spike trains (MUCST) were used to calculate intramuscular, intermuscular, and Force-MUCST coherence in the delta (0-5 Hz), alpha (5-15 Hz), and beta (15-35 Hz) frequency bands. Changing the ankle position from a shortened to lengthened position resulted in increased GM-SM coherence in the delta and alpha bands and improved force steadiness, while intramuscular coherence and Force-MUCST coherence decreased in the alpha band. However, there were minimal changes in beta band coherence. Motor unit recruitment thresholds of GM and SL were reduced with muscle lengthening, while GL showed greater thresholds at lengthened positions, and SM was unaffected by ankle position. These findings highlight the role of inhibitory inputs associated with changes in muscle length and the modulation of these inputs by neighboring synergistic muscles. This study reveals a neuromuscular control strategy that modulates common synaptic inputs and motor unit recruitment of triceps surae to maintain force output during isometric plantarflexions at varying muscle lengths.

**Key points:**

- When comparing isometric contractions at different ankle joint angles, a lengthened position resulted in greater shared common inputs between the medial head of the gastrocnemius and the medial compartment of the soleus in delta and alpha frequency bands during submaximal force production.
- Increased muscle length resulted in improved force steadiness, as well as reduced common synaptic inputs within each triceps surae muscle and coupling between force output and neural drive in the alpha band.
- Shifting the ankle position from a shortened to a lengthened position resulted in a reduced motor unit recruitment threshold of GM and SL. However, GL exhibited an opposite pattern, showing higher recruitment thresholds at lengthened positions, while SM was not affected by changes in muscle length.
- This study reveals a neuromuscular control strategy that integrates afferent feedback and muscle mechanical properties while modulating common descending inputs to synergistic plantarflexors.

## Introduction

Triceps surae muscles, which include the gastrocnemius and soleus, share a common insertion and function as synergists for ankle plantarflexion (Dalmau-Pastor et al., 2014). Despite this shared function, they exhibit distinct characteristics, including differences in muscle fiber types (Johnson et al., 1973) and architectural features like fascicle length and angle (Crouzier et al., 2018; Kawakami et al., 1998). Furthermore, changes in ankle position can alter force-length relationships of triceps surae muscles, potentially influencing force-sharing strategies (Kawakami et al., 1998). Our prior research identified two primary common inputs driving triceps surae motoneurons, with these inputs distributed consistently across ankle angles (Levine et al., 2023). Common synaptic inputs to motoneurons may arise from descending, spinal interneuronal and sensory pathways (Hug et al., 2023). A frequency-domain analysis that quantifies coherence between motor unit spike trains in different frequency bands can reveal the spectral properties and origins of these inputs (Brown, 2000; Grosse et al., 2002). Previous studies have shown minimal delta-band coherence—representing effective neural drive (Farina et al., 2014) —between gastrocnemius medialis (GM) and lateralis (GL) (Hug et al., 2021; Rossato et al., 2022), allowing flexible motor control at the ankle (Rossato et al., 2022). In contrast, the medial (SM) and lateral soleus (SL) compartments share strong common inputs, comparable to those within each compartment (Aeles et al., 2023). Importantly, cadaver and imaging studies have shown that the medial portions of gastrocnemius (Crouzier et al., 2018) and soleus (Kimura et al., 2021) have greater physiological cross-sectional area and muscle volume than their lateral counterparts, suggesting greater force-generating capacity (Crouzier et al., 2018). However, shared common inputs in GM-SM and GL-SL muscle pairs remain unexplored. Furthermore, most studies on the effects of altering ankle position have focused solely on the delta band within and across these synergists, leaving the role of peripheral (alpha band) and corticospinal (beta band) common inputs unclear.

Oscillations of common synaptic inputs to motoneurons in the alpha band include the central elements of physiological tremor (Conway et al., 1995). Furthermore, alpha band coherence is affected by afferent feedback (Laine et al., 2021) and has been linked to force steadiness (Budini et al., 2014; Laine et al., 2013), which can be estimated by quantifying coupling between force output and neural drive in low-frequency bands (Laine et al., 2016; Negro et al., 2009). Changes in muscle length induce modulations such as gamma fusimotor drive (Ellaway et al., 2015; Jalaleddini et al., 2017) and muscle mechanical properties (Cabral et al., 2024), which in turn affect oscillations in common synaptic inputs to motoneurons through peripheral afferent feedback. Importantly, in addition to shared descending drive across triceps surae muscles, neighboring synergistic muscles typically show coordinated afferent activity and reflex responses (Laine et al., 2016). Furthermore, the afferent feedback may also influence sensorimotor integration and recalibrate descending corticospinal inputs (Witham et al., 2011), which can be estimated by motor unit coherence in the beta band (Brown, 2000; McManus et al., 2019). Therefore, understanding how altering ankle position modulates common synaptic inputs within and across triceps surae muscles in the alpha and beta frequency bands will reveal the neuromuscular mechanism underlying force-length relationship and force sharing strategy of triceps surae muscles.

The force generated by a muscle during voluntary contraction is determined by both the number of motor units recruited and their discharge rates (Enoka & Duchateau, 2017). Our prior study showed that discharge rates of GL and Soleus (SL and SM combined) are significantly greater at shortened compared to lengthened position, while discharge rates of GM remain unchanged (Levine et al., 2023). According to the size principle (Henneman, 1957) and prior research (Héroux et al., 2014), the soleus has lower motor unit recruitment thresholds than gastrocnemius due to their muscle fiber composition: Soleus is predominantly slow-twitch while gastrocnemius consists of approximately 50% fast-twitch and 50% slow-twitch muscle fibers (Johnson et al., 1973). Furthermore, muscle fiber lengths and pennation angles are similar for GM and soleus (40 mm and 20°), but different from those of GL (80 mm and 10°) (Kawakami et al., 1998; Maganaris et al., 1998). This may explain previous findings that the motor unit recruitment threshold of GL is significantly higher than that of GM and soleus during standing and voluntary plantarflexion, suggesting that GM and GL may function within different ankle ranges during plantarflexion (Héroux et al., 2014). Previous studies demonstrated greater motor unit recruitment threshold of GM at shortened position compared to lengthened position (Kennedy & Cresswell, 2001; Nishimura & Nakajima, 2002). However, it remains unknown how altering muscle length modulates motor unit recruitment threshold of GL and soleus to produce force.

Therefore, the aim of this study was to investigate how altering muscle length, by changing ankle position, affects common synaptic inputs within and across the triceps surae muscles in the delta, alpha and beta frequency bands, as well as force control and motor unit recruitment. Given that alpha band coherence is highly affected by peripheral afferent feedback (Laine et al., 2021) and that reduced intramuscular coherence has been observed in the TA as muscle length increases (Cabral et al., 2024), we hypothesized that increasing muscle length will result in decreased intramuscular and intermuscular alpha band coherence. Since the distribution of common drive to motoneurons in the triceps surae remains robust across different ankle angles (Levine et al., 2023) and beta band coherence does not directly contribute to voluntary force (Zicher et al., 2023), we hypothesize minimal changes in delta and beta band intramuscular coherence across ankle positions. As our previous study on the same dataset observed the lowest motor unit discharge rates at the lengthened position (Levine et al., 2023) and Force-EMG coherence reflects the patterns of force fluctuations (Laine et al., 2016), we hypothesized that force fluctuations and Force-MUCST coherence will be reduced as muscle lengthens. Based on the differences in muscle architecture and fiber composition between GM/soleus and GL (Héroux et al., 2014; Kennedy & Cresswell, 2001; Nishimura & Nakajima, 2002), we hypothesize that the motor unit recruitment threshold of GM and soleus will decrease as muscle length increases, while GL will exhibit an opposite pattern.

## Methods

### Participants

Sixteen healthy young adults (8 females, 8 males, age: 28.4 ± 7.9 years, BMI: 23.6 ± 2.7) participated in the study. They had no history of knee or ankle pain for the past six months, and did not experience any pain in the lower leg that could affect voluntary contractions. All participants provided their informed written consent prior to participating, and all experimental procedures were approved by the Institutional Review Board of the Northwestern University (IRB No. STU00212191). The same dataset has been used in our previous work investigating motor neuron synergies (Levine et al., 2023).

### Experimental set up

Participants were seated on an isokinetic dynamometer (Biodex System 4 Pro; Biodex Medical, Shirley, NY, USA) with their upper leg, waist and both shoulders fixed by straps. They sat with their hips flexed at a 90° angle, knee fully extended, and the foot of the dominant leg (i.e. right leg for all the participants) secured to a pedal (Figure. 1).

**Figure 1.**
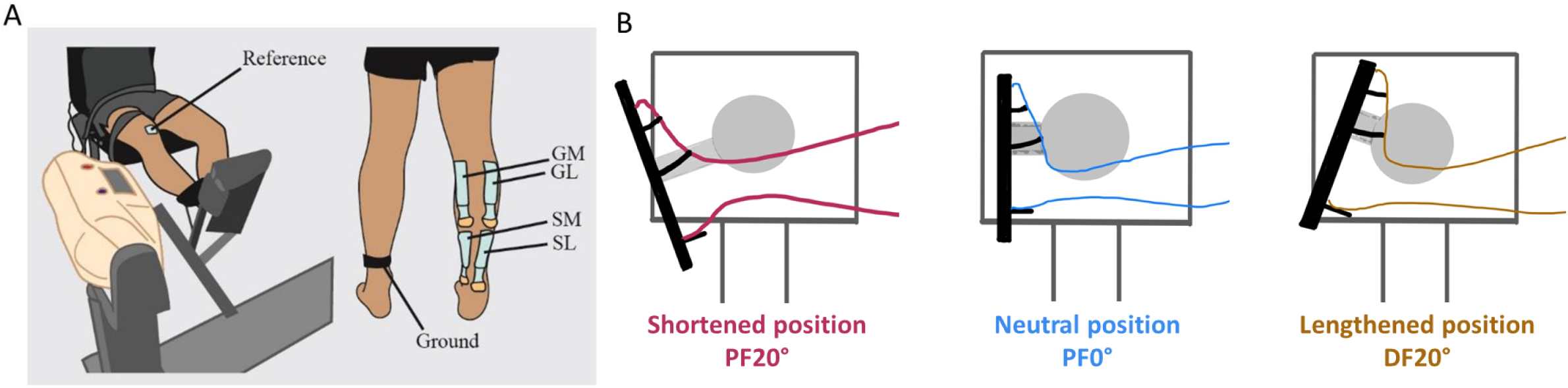
HDsEMG electrodes placement (A) and experimental setup (B).

Participants’ skin was shaved, scrubbed, and any dead skin was removed using wet towels. Four high density surface electromyography (HDsEMG) grids, each containing 64 electrodes (GR08MM1305, 13 × 5 gold-coated electrodes with one electrode absent on a corner; interelectrode distance, 8 mm; OT Bioelettronica, Italy), were placed over medial and lateral compartments of soleus (SM and SL), and medial and lateral heads of gastrocnemius (GM and GL) (Figure 1). Each grid was prepared by applying a semi-disposable bi-adhesive foam layer (FOA08MM1305, OT Bioelettronica, Italy) to hold the grid of electrodes to the skin. The cavities of the adhesive layer were equally filled with conductive cream (AC Cream, Spes Medica, Genoa, Italy) to ensure skin–electrode contact. A reference electrode (Ag/AgCl snap electrodes; Noraxon, USA) was placed over the patella. Two wet electrode bands were placed around the ankles of both legs to serve as ground electrodes. The EMG signals were recorded in monopolar mode, band-pass filtered at 10-500Hz, and digitized at a sampling rate of 2048 Hz using a multichannel EMG system (EMG-Quattrocento, OTBioelettronica, Italy). Signals were recorded using OT Biolab+ software (OT Bioelettronica, Italy).

### Experimental procedure

After electrode placement, participants performed several practice contractions. They were instructed to perform a standardized warm-up by pushing against the pedal with gradually increasing force, reaching approximately 50% of their perceived maximal effort, while keeping their ankle in a neutral position (0°). Following three contractions at this level, participants were then instructed to gradually increase the force to approximately 80-90% of their perceived maximal effort for one contraction.

The experimental protocol included isometric ankle plantarflexions at 20% of maximum voluntary contraction (MVC). Three ankle positions were evaluated: the ankle plantarflexed at 20° (PF20°), at the neural position (PF0°) and dorsiflexed at 20° (DF20°). Three different conditions were assessed in a randomized order. For each condition, participants were first instructed to rapidly increase the isometric plantarflexion force exerted by their ankle from baseline to maximum and hold it at maximum for 5 seconds. They performed two MVCs with 1-minute rest intervals between each while receiving verbal encouragement. The peak MVC was calculated as the average maximal torque of the contractions and was used to set the 20% MVC target torque. After resting for 1 minute, participants performed three trapezoidal contractions, each comprising a 15s linear ramp-up, a 30s plateau, and a 15s linear ramp-down. There was a 10s resting time between each trapezoidal contraction. Visual feedback of the target and torque output was displayed on a monitor in front of the participants. Torque signals were low-pass filtered at 20 Hz using a third order Butterworth filter, and were baseline corrected to eliminate the weight of the testing leg and the pedal.

### Data analysis

MATLAB custom-written scripts (R2023b; The MathWorks, Natick, MA, USA) were used for data analysis.

#### Force output

To assess the force fluctuations, the 30-second plateau of the trapezoidal contractions was selected to calculate the coefficient of variation (CV) of force, a measure of force steadiness. To assess physiological tremor, the power spectral density of the torque in the alpha band was calculated using 1 s Hanning windows with 1948 samples of overlap (“pwelch” function in MATLAB) (Cabral et al., 2024).

#### EMG decomposition

Channels with low signal-to-noise ratio or artifacts were discarded by visual inspection. The EMG signals were then decomposed into motor unit pulse trains with a blind source separation algorithm (Negro et al., 2016) using DEMUSE software (A. Frančič & A. Holobar, 2021). All decomposed spike trains were manually edited for false negatives and false positives by one experienced investigator and double-checked by another experienced investigator, and duplicates were removed, following previously published guidelines (Del Vecchio et al., 2020). Motor units with a pulse-to-noise ratio >30 dB were retained for further analysis (Holobar et al., 2014).

#### Motor unit recruitment threshold

Motor unit recruitment threshold was defined as the torque at which the motor unit began to discharge during each isometric ramp contraction (Martinez-Valdes et al., 2022). Recruitment threshold was then expressed as a percentage of the MVC torque obtained at the same ankle angle.

#### Coherence

Only motor units continuously firing during the 30-s force plateau window were included in the coherence analysis. Intramuscular coherence within each triceps surae muscle was calculated using two cumulative spike trains (CSTs) of equal size. Each CST was composed of the summed discharge rates of three motor units, randomly selected from the manually edited motor units in one muscle. Therefore, the minimum number of motor units required from one muscle was six for calculating intramuscular coherence. One subject was excluded from this analysis because the number of analyzable motor units from any of the four muscles was less than six.

Intermuscular coherence was calculated using two cumulative spike trains (CSTs) of equal size from two different muscles. Each CST consisted of the summed discharge rates of three motor units, randomly selected from the manually edited motor units in each muscle. A minimum of three motor units from each muscle was required to calculate intermuscular coherence. One subject was excluded from this analysis because the number of analyzable motor units in any of the four muscles was less than three. To assess the coupling between force output and neural drive, coherence was calculated between force and CSTs of all motor units (MUs) from each muscle (Cabral et al., 2024). Since the number of motor units included in the analysis can affect coherence values (Farina & Negro, 2015), we included the number of motor units as a covariate in the statistical analysis. Force-MUCST coherence was only assessed in the delta and alpha frequency bands because low-frequency oscillations in motoneuron activity have the greatest impact on force generation and fluctuations due to the low-pass filtering effect of neural drive on muscle force (Negro et al., 2009). Additionally, oscillations in the beta band do not contribute directly to force output (Zicher et al., 2023).

For the 30-second plateau of each trapezoidal contraction, coherence analysis was performed on the pooled CSTs using random combinations from 100 iterations. Using “mscohere” function in MATLAB, coherence was estimated with Welch’s averaged periodogram with a non-overlapping Hanning window of 1 s duration (Del Vecchio et al., 2019; Rossato et al., 2022). The average coherence of these 100 iterations was then compared to the level for significant coherence (confidence limit, CL), as determined using Eq. 1, where L is the total number of segments.

CL = 1- 0.05^ (1/(L −1) (Eq. 1, Rosenberg et al., 1989)

Coherence values greater than CL were then z-transformed using Eq. 2:

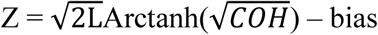 (Eq. 2, Del Vecchio et al., 2019)

Bias was defined for each participant as the mean value of z-scores between 250 and 500 Hz, as no significant coherence is expected in this frequency range (Cabral et al., 2024; Del Vecchio et al., 2019). Mean z-transformed pooled coherence at frequency ranges of 0-5 Hz (delta band), 5-15 Hz (alpha band), and 15-35 Hz (beta band) (Brown, 2000) was calculated for further statistical analysis. Figure 2 shows the pooled coherence across all participants for a representative muscle pair at three different ankle positions.

**Figure 2.**
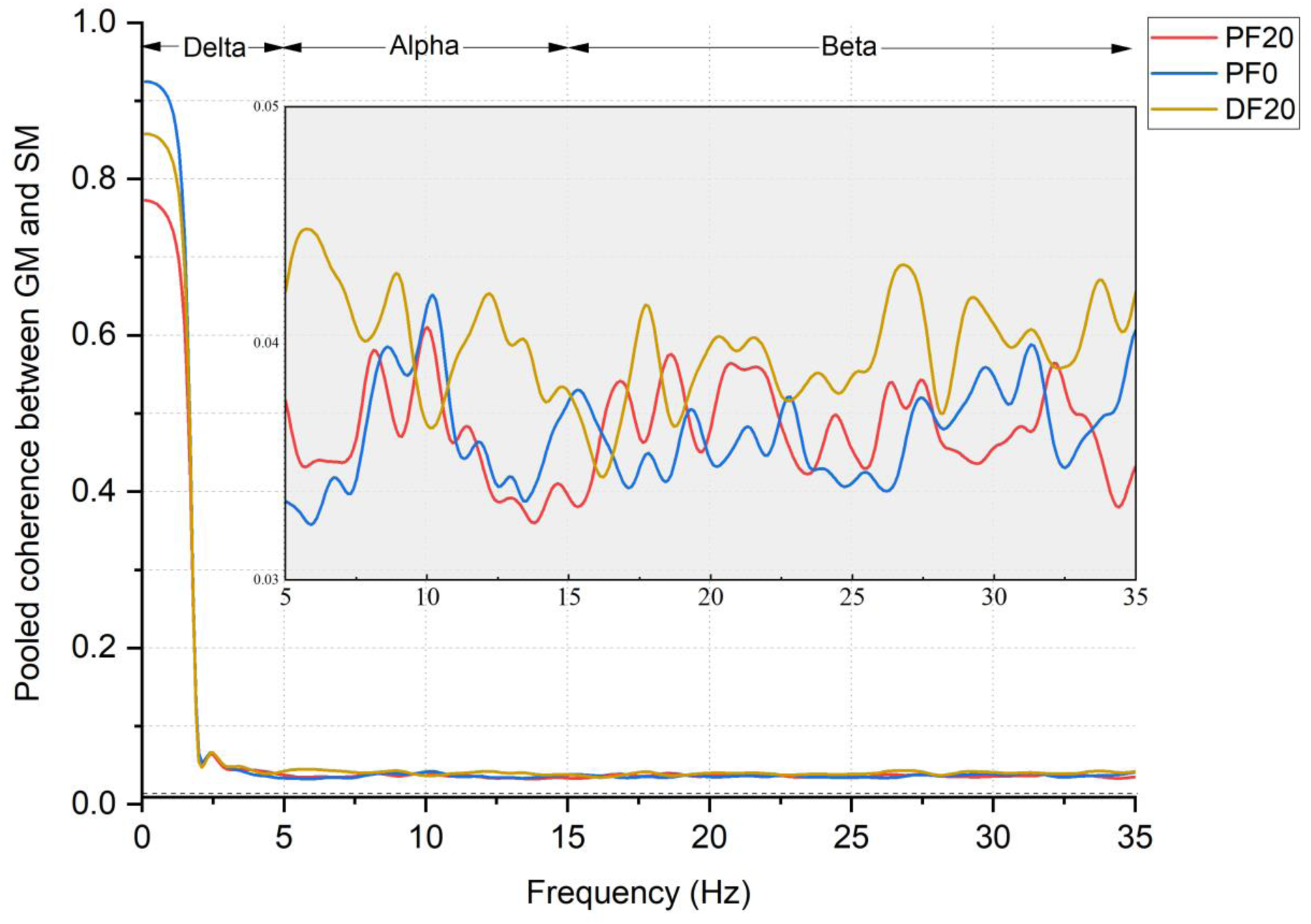
Pooled GM-SM coherence across all participants at three ankle positions. The dashed line represents the confidence level.

### Statistical analysis

SPSS 26.0.0.1 was used for statistical analysis. A Shapiro–Wilk’s test was used to test for normal distribution of the data. A linear mixed model with a fixed effect of Position (PF20°, PF0°, DF20°) was applied to assess MVC torque, CV of force output, and power spectral density of force in the alpha band, with participant and contraction number included as random effects. A linear mixed model with a fixed effect of Position (PF20°, PF0°, DF20°) and Muscle (GM, GL, SM, SL) was applied to evaluate motor unit recruitment threshold, with participant and contraction number included as random effects. Mean z-transformed coherence over the 30s plateau in the delta, alpha, and beta bands were fit with a linear mixed model with fixed effects of Muscle (GM, GL, SM, SL, GM-GL, SM-SL, GL-SL, GM-SM), Position (PF20, PF0, DF20) and their interaction, with participant and contraction number as random effects (McManus et al., 2019). For the analysis of Force-MUCST coherence, number of motor units was included as a covariate. Statistical significance was set at *p* < 0.05 for all analyses, and data are presented as mean ± standard error.

## Results

### MVC torque

MVC torque at PF0° (737.42 ± 356.10 Nm) was similar to that at DF20° (807.44 ± 458.30 Nm), and both were significantly greater than MVC torque at PF20° (396.45 ± 226.92 Nm) (both *p* = 0.000).

### Force fluctuations

There was a main effect of Position [F (2, 116) = 22.67, *p* = 0.000] on the CV of force output. CV of force was greater at PF20° (1.34 ± 0.08) compared to that at PF0° (1.22 ± 0.08, *p* = 0.016) and DF20° (1.01 ± 0.08, *p* = 0.000), and it was greater at PF0° than at DF20° (*p* = 0.000). Therefore, force fluctuations decreased as muscle length increased (Figure 3A).

**Figure 3.**
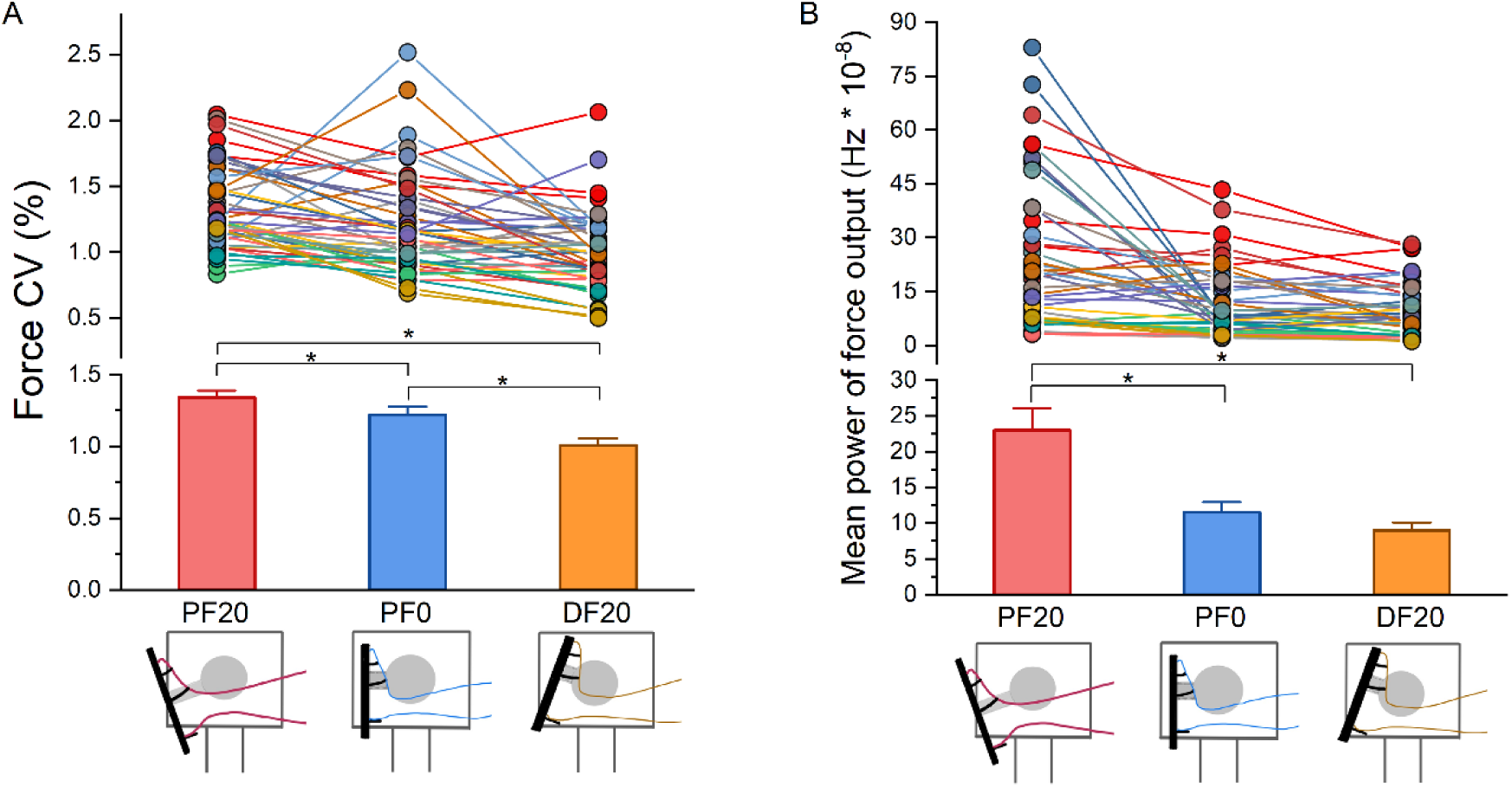
CV (A) and mean power (B) of force output.

There was a main effect of Position [F (2, 111) = 25.93, *p* = 0.000] on mean power of force output in the alpha band. Mean power of force output in the alpha band at PF20° (23.92 ± 3.20) was greater than that at PF0° (11.467 ± 3.15, *p* = 0.000) and DF20° (8.74 ± 3.20, *p* = 0.000), but no differences were observed between PF0° and DF20° (Figure 3B).

### Number of Decomposed Motor Units

There were 1,665 motor units manually edited and screened for continuous firing for further analysis. Table 1 shows the total number of decomposed motor units per muscle and condition. Motor units were not tracked across angles to maximize the number of units available for coherence analysis.

**Table 1.** Number of motor units used for analysis

| No. of MUs | GM | GL | SL | SM | Total |
| --- | --- | --- | --- | --- | --- |
| PF20° | 172 | 139 | 151 | 134 | 596 |
| PF0° | 166 | 138 | 153 | 137 | 594 |
| DF20° | 162 | 94 | 122 | 97 | 475 |
| Total | 500 | 371 | 426 | 368 | 1665 |

### Delta band (0-5 Hz)

#### Intramuscular and intermuscular coherence

There was a main effect of Muscle [F (7, 681) = 37.87, *p* = 0.000], a main effect of Position [F (2, 679) = 5.73, *p* = 0.003], and a Muscle x Position interaction [F (14, 678) = 1.74, *p* = 0.044] on delta band motor unit coherence. Coherence within SL was greater at DF20° compared to PF20° (*p* = 0.033) and PF0° (*p* = 0.008), and coherence within SM was greater at PF0° than PF20° (*p* = 0.032). However, delta-band coherence within GM and GL did not change with position. Intermuscular coherence between GM and SM was lower at PF20° compared to DF20° (*p* = 0.001) and PF0° (*p* = 0.007). Altering ankle position did not affect intermuscular coherence in the other three muscle pairs (Figure 4).

**Figure 4.**
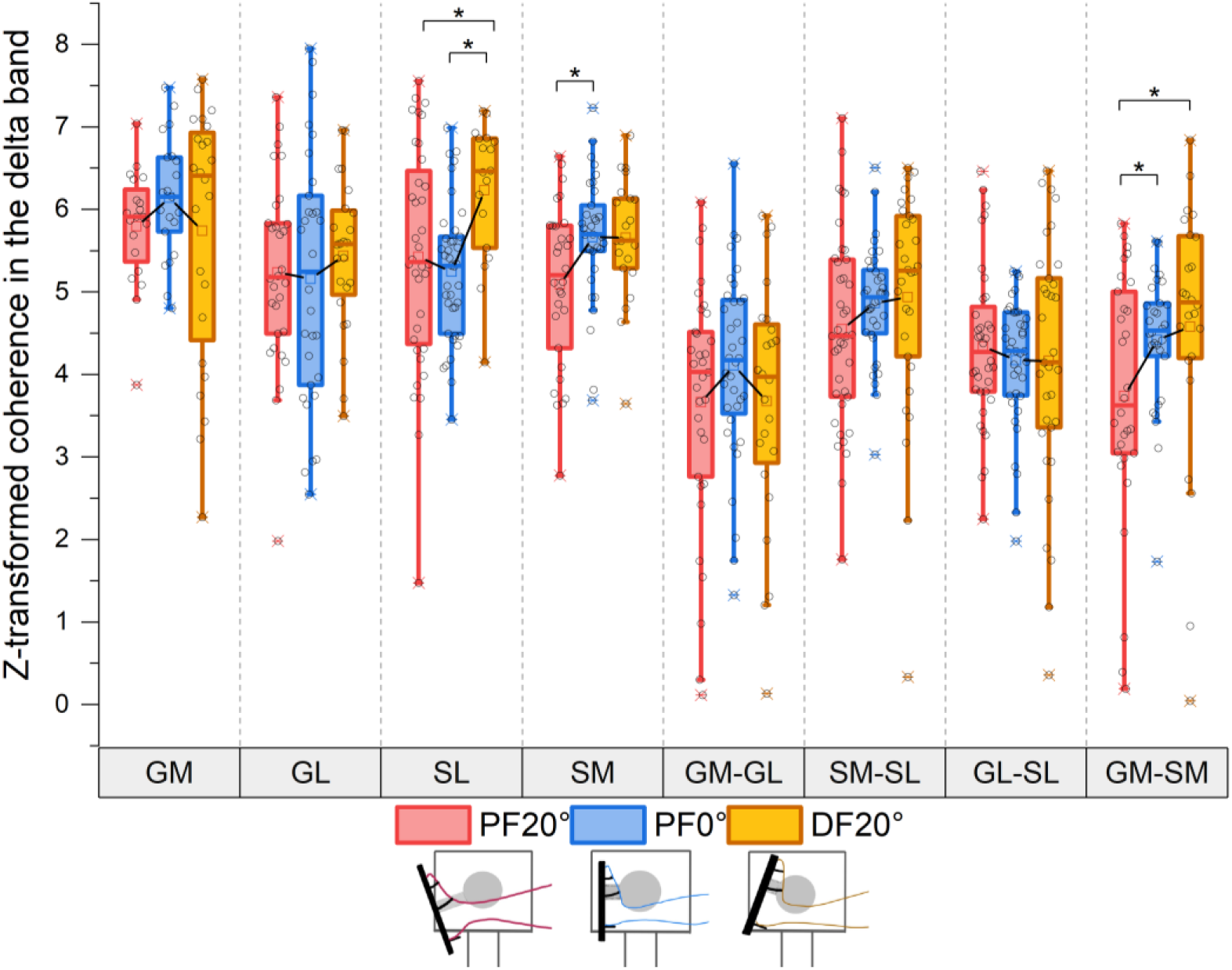
Mean z-transformed intramuscular and intermuscular coherence in the delta band. *Statistical significance is reported for *p* < 0.05.

#### Force-MUCST coherence

No significant main effects or interactions were found on delta band coherence between force output and MUCSTs (main effect of Muscle: *p* = 0.598, main effect of Position: *p* = 0.326, Muscle x Position interaction: *p* = 0.896).

### Alpha band

#### Intramuscular and intermuscular coherence

There was a main effect of Muscle [F (7, 534) = 27.75, *p* = 0.000], a main effect of Position [F (2, 534) = 11.31, *p* = 0.000], and a Muscle x Position interaction [F (14, 533) = 1.99, *p* = 0.017] on alpha band motor unit coherence. Coherence within GM at PF20° and PF0° were greater than that at DF20° (both *p* = 0.000). Coherence within GL was greater at PF20° than at PF0° (*p* = 0.022), and coherence within SM at PF20° was greater than PF0° (*p* = 0.023) and DF20° (*p* = 0.019). However, coherence within SL did not change with position. Intermuscular coherence in GM-GL at PF20° was greater than that at PF0° (*p* = 0.043). GM-SM coherence at PF0° was increased compared to DF20° (*p* = 0.033), but other muscle pairs showed no change of coherence with ankle position (Figure 5).

**Figure 5.**
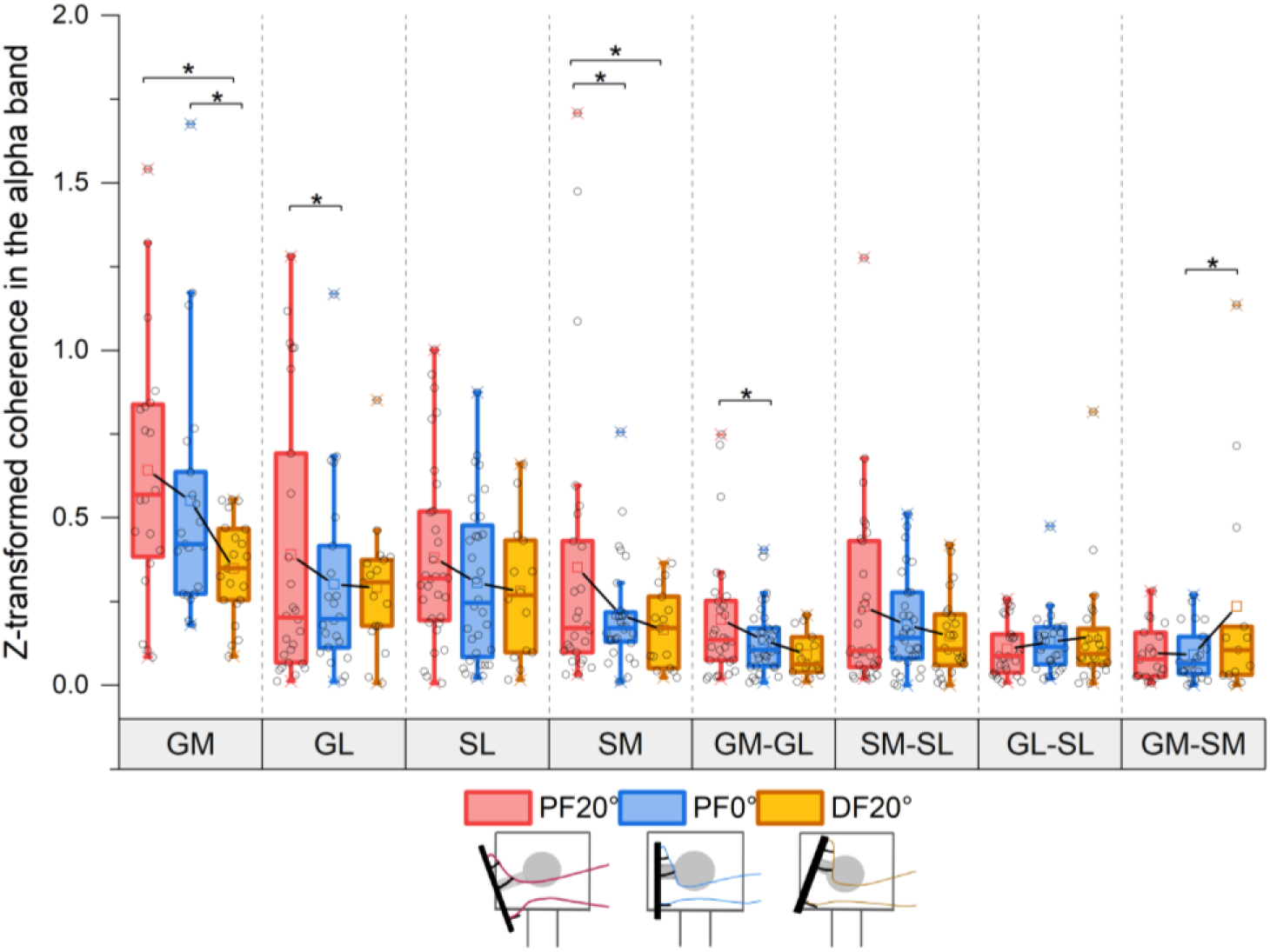
Mean z-transformed intramuscular and intermuscular coherence in the alpha band. *Statistical significance is reported for *p* < 0.05.

#### Force-MUCST coherence

There were a main effect of Position [F (2, 130) = 10.06, *p* = 0.000] on alpha band coherence between force output and MUCSTs. No main effect of Muscle [F (3, 130) = 2.658, *p* = 0.051] nor Position x Muscle interaction [F (6, 130) = 0.77 *p* = 0.592] were found. Force-MUCST at PF20° was greater than that at PF0° (*p* = 0.026) and DF20° (*p* = 0.000). Force-MUCST at PF0° was greater than that at DF0° (*p* = 0.028) (Figure 6).

**Figure 6.**
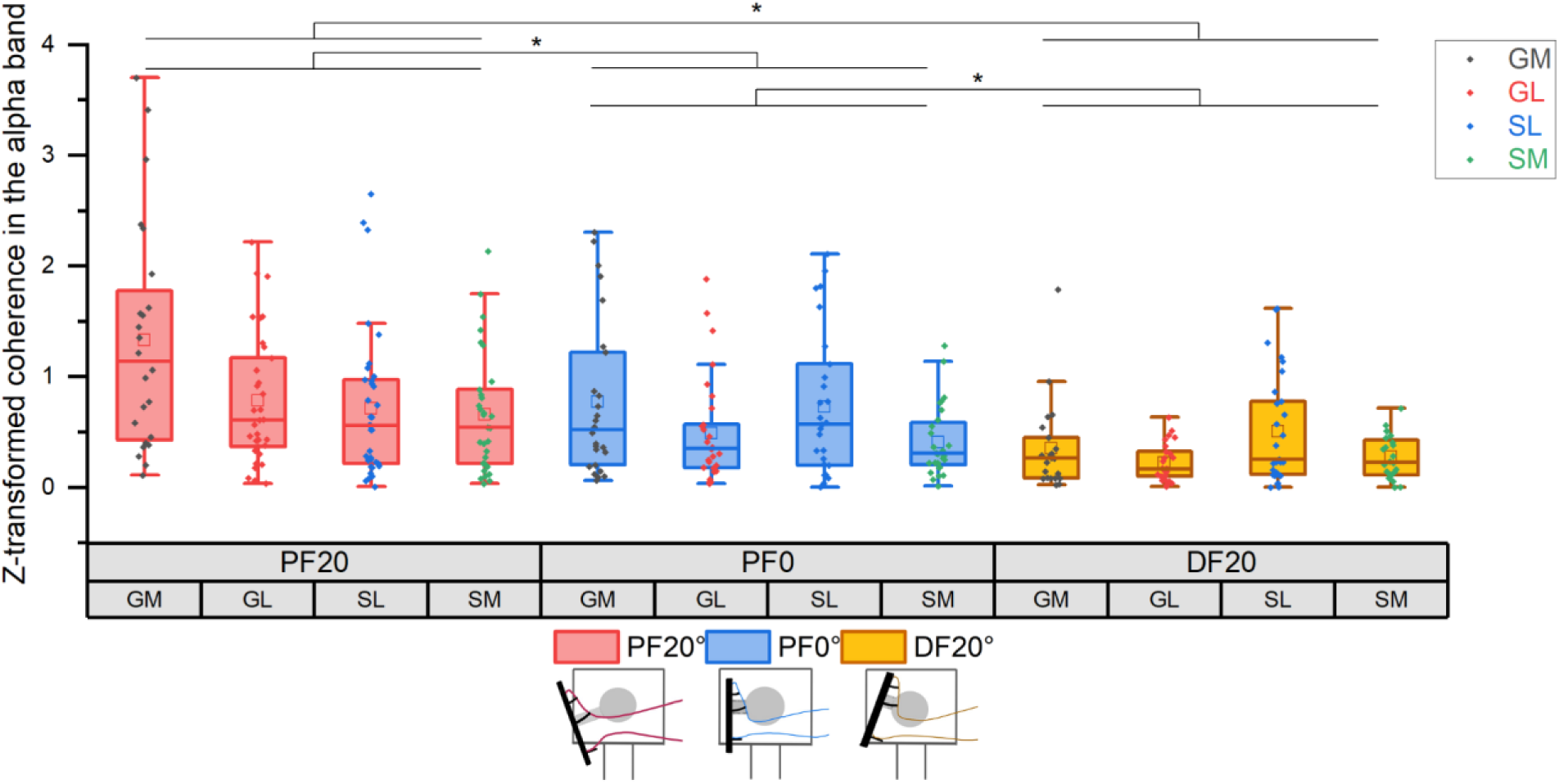
Mean z-transformed Force-MUCST coherence in the alpha band. *Statistical significance is reported for *p* < 0.05.

### Beta band

There was a main effect of Muscle [F (7, 489) = 33.42, *p* = 0.000], but no significant main effect of Position [F (2, 489 = 0.97, *p* = 0.382] nor Muscle x Position interaction [F (14, 488) = 0.89, *p* = 0.571] on beta band coherence. When pooled across the three positions, beta band coherence within GM was greater than in other muscles and muscle pairs (all *p* = 0.000). Coherence within GL (all *p* = 0.000) and within SL (all *p* < 0.01) were greater than that within SM and between all muscle pairs. Intermuscular coherence in the beta band did not differ among muscle pairs (Figure 7).

**Figure 7.**
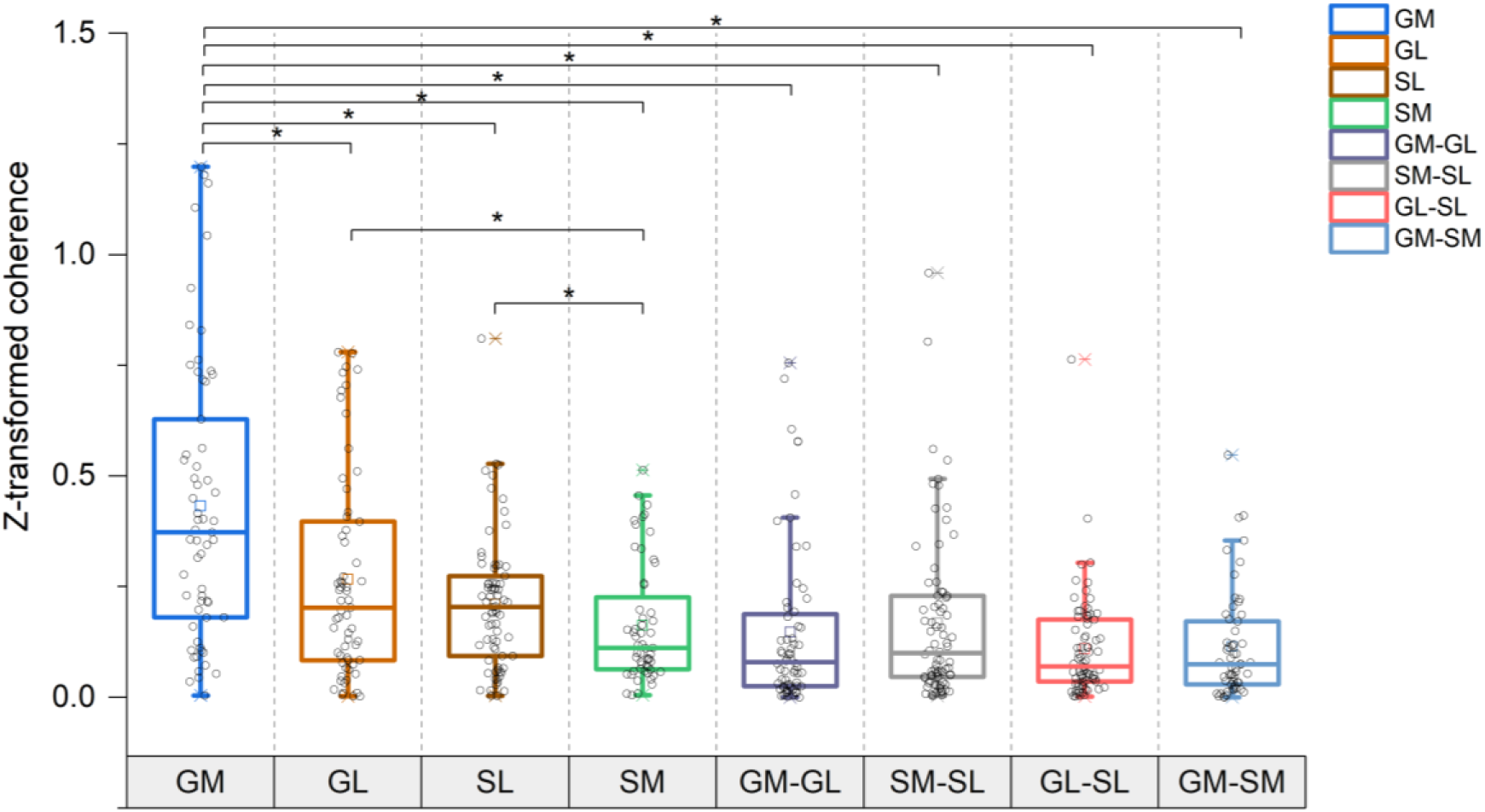
Mean z-transformed intramuscular and intermuscular coherence in the beta band pooled across three ankle positions. *Statistical significance is reported for *p* < 0.05.

### Motor unit recruitment threshold

There was a main effect of muscle [F (3, 1634) = 17.40, *p* = 0.000], a main effect of Position [F (2, 1647) = 7.40, *p* = 0.001], and a Muscle x Position interaction [F (6, 1644) = 5.01, *p* = 0.000] on motor unit recruitment threshold. When pooled across position, recruitment threshold of GL motor units was greater (recruited at greater forces) than that of GM, while both were greater than that of SL and SM (both *p* = 0.000). Recruitment threshold of GM motor units decreased as ankle position shifted from PF20° to DF20° (all *p* < 0.05), while recruitment threshold of GL motor units at PF20° was lower than that at PF0° and DF20° (both *p* < 0.05). Recruitment threshold of SL motor units at DF20° was lower than that at PF0° and PF20° (both *p* ≤ 0.01). Recruitment threshold of SM motor units remained unchanged across ankle positions (Figure. 8A).

**Figure 8.**
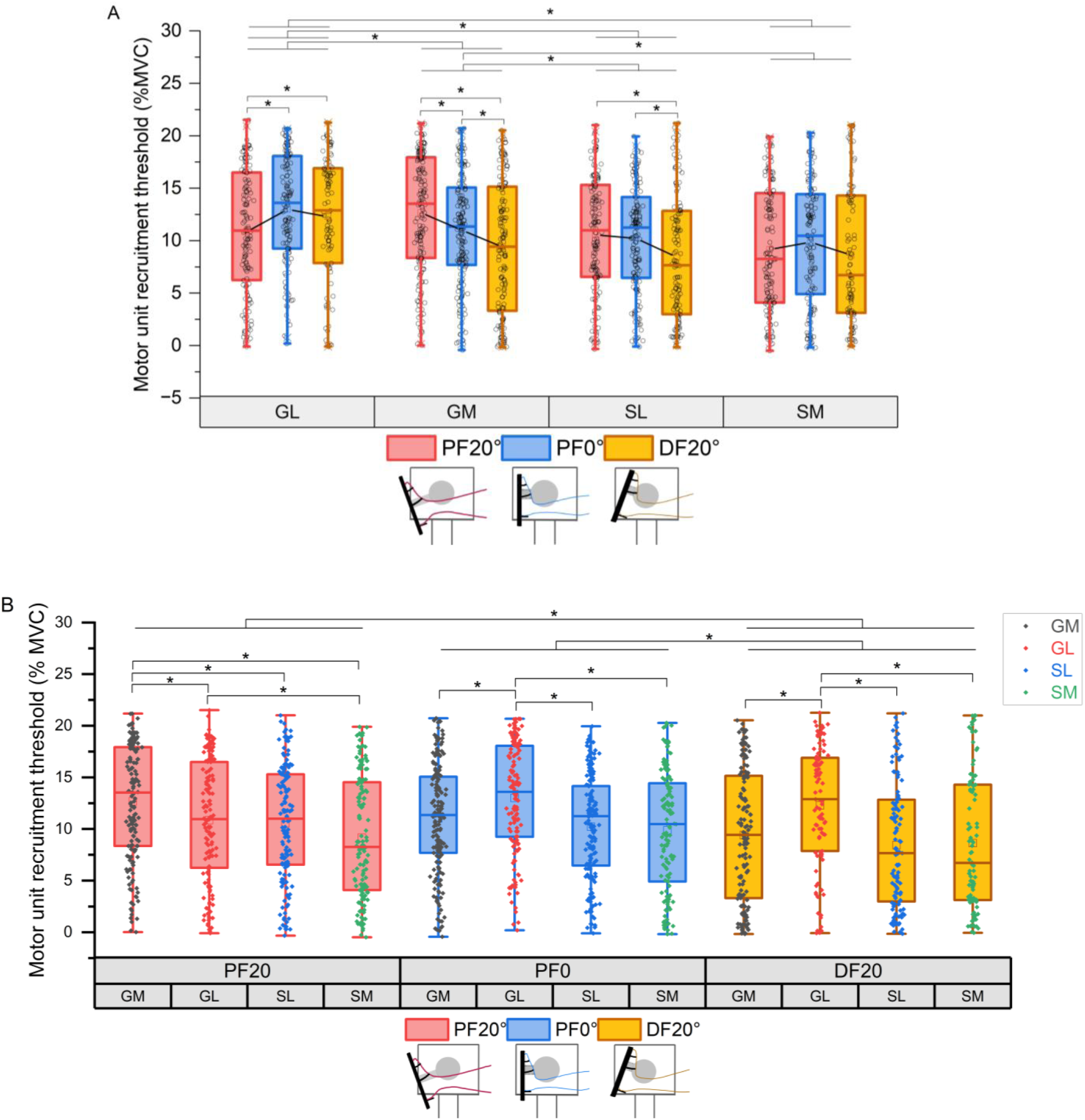
Motor unit recruitment threshold normalized to maximal torque of each triceps surae muscle (A) at three ankle positions (B). *Statistical significance is reported for *p* < 0.05.

Recruitment threshold at DF20° was lower than that at neutral (*p* = 0.000) and PF20° (*p* = 0.004), regardless of muscle. At PF20°, recruitment threshold of GM motor units was greater than other three muscles (all *p* < 0.01), and recruitment threshold of GL motor units was greater than that of SM (*p* = 0.035). At PF0°, GM, SL, SM showed similar recruitment threshold, but lower than that of GL (all *p* < 0.01). At DF20°, recruitment threshold of GL motor units was greater than that of other three muscles (all *p* = 0.000) (Figure. 8B).

## Discussion

Changing the ankle joint angle from a shortened (plantarflexed) to lengthened (dorsiflexed) position resulted in increased shared synaptic inputs (i.e. motor unit coherence) between the medial head of the gastrocnemius and the medial compartment of the soleus muscle (GM-SM) but not between other muscle pairs, particularly in the delta and alpha frequency bands. Meanwhile, muscle lengthening results in decreased common synaptic inputs within each triceps surae muscle and coupling between force output and neural drive (i.e. Force-MUCST coherence) in the alpha band as well as improved plantarflexion force steadiness. However, altering ankle joint angle had minimal impact on the shared corticospinal inputs (i.e. beta band coherence) across these muscles. This study reveals a neuromuscular control strategy that modulates common synaptic inputs, force control and motor unit recruitment of triceps surae during isometric plantarflexions at varying muscle lengths.

### Enhanced neural coupling across medial triceps surae muscles with muscle lengthening

We found GM-SM coherence in the delta and alpha frequency bands increased with increasing muscle length. Delta band coherence is widely used to estimate common synaptic inputs to motoneuron pools that produce muscle force (Farina et al., 2014; Farina & Negro, 2015; Hug et al., 2021; Laine et al., 2015; Rossato et al., 2022). Oscillations in the delta band can be found in muscles with and without muscle spindles (Kamen & De Luca, 1992), suggesting that delta band motor unit coherence may not depend on the presence of muscle spindles. A previous study reported significant delta band coherence during pedaling between synergist muscles responsible for ankle stabilization, primarily driven by the coordination between the GM and soleus muscles (De Marchis et al., 2015). In addition, greater intermuscular alpha band coherence has been demonstrated between muscles with higher weights within a synergy (Laine et al., 2021; Ortega-Auriol et al., 2023). Therefore, our findings suggest that increased muscle length during isometric plantarflexion leads to greater shared common inputs across the medial compartments of the gastrocnemius and soleus. Since the medial portions of gastrocnemius (Crouzier et al., 2018) and soleus (Kimura et al., 2021) have greater physiological cross-sectional area and muscle volume than their lateral portions, this neuromuscular strategy enhances neural coupling in the compartments with greater force-generating capacity, helping to maintain force output more effectively. This change of GM-SM intermuscular coherence also supports the two motor neuron synergies hypothesis we proposed in our previous work (Levine et al., 2023), as there is clear evidence of an increase in coupling of neural drive in the medial compartments.

### Reduced alpha band intramuscular coherence, physiological tremor and force fluctuations as muscle lengthened

The present study found that shifting the ankle joint from a shortened to lengthened position led to reduced force CV and physiological tremor (i.e. mean power) of force output during isometric plantarflexion. Interestingly, we also observed reduced intramuscular coherence and Force-MUCST coherence within GM, GL and SM in the alpha band. These results suggest that at lengthened position compared to shortened position, there were enhanced force steadiness, reduced neural drive within each muscle, and decreased coupling between neural drive and force output in the alpha band. Alpha band oscillations include the central elements of physiological tremor (Conway et al., 1995). Reduced intramuscular alpha band coherence and force fluctuations at lengthened position have been demonstrated in previous research (Cabral et al., 2024; Jalaleddini et al., 2017) and can be attributed to the following modulations as muscle length increases: 1) Reduction in gamma static fusimotor drive (Ellaway et al., 2015; Jalaleddini et al., 2017) induced by activities of secondary endings in the muscle spindle, which are more associated with postural control and static limb positioning (Banks et al., 2021). This altered activity of muscle spindles is influenced not only by length changes in the muscle where they are located but also by length changes in co-activated muscles (Laine et al., 2016; Smilde et al., 2016); 2) Decreased pre-contraction muscle tendon unit tension, resulting in altered peripheral contractile properties of motor units (Cabral et al., 2024); 3) Changes in reciprocal inhibition from the antagonist muscle (i.e. tibialis anterior) during plantarflexion (Yavuz et al., 2018); 4) Increased recurrent inhibition to motor neurons from Renshaw cells (Dideriksen et al., 2015), which has been shown to reduce physiological tremor (Edgley et al., 2021; Williams & Baker, 2009).

### Reduced coupling between neural drive and force output in the alpha band as muscle lengthened

The present study evaluated Force- MUCST coherence in delta and alpha frequency bands, which provide an estimate of the amount that a muscle contributes to the net force output (Enoka & Farina, 2021). We found reduced coherence between force output and MUCST in the alpha band as a result of muscle lengthening, which aligns with previous research that demonstrated a similar pattern of Force-EMG coupling to the behavior of force fluctuations (Laine et al., 2016). However, Cabral et al., (2024) observed increased Force-MUCST alpha band coherence in tibialis anterior during isometric dorsiflexion at lengthened position compared to shortened position, which is opposite of our observations. Furthermore, Cabral et al. found that changes in muscle length of the tibialis anterior do not affect force CV, which may be explained by the unchanged motor unit discharge rate in tibialis anterior across muscle lengths during submaximal isometric dorsiflexion (Cabral et al., 2024; Tsatsaki et al., 2022). However, our prior work on the same dataset found that discharge rates of GL and Soleus (SL and SM combined), but not GM, are indeed significantly greater at shortened compared to lengthened position (Levine et al., 2023). Different factors may account for the variations in force steadiness between dorsiflexion and plantarflexion, as well as differences in motor unit discharge rates between the tibialis anterior and the triceps surae muscles. The strength of reciprocal inhibition in the tibialis anterior motor units is fourfold greater than that of the GM and the soleus motor units (Yavuz et al., 2018), which may primarily be due to the fact that the muscle spindle density of the triceps surae is double that of the tibialis anterior (De Luca & Kline, 2012). Additionally, afferent activity and reflex responses from neighboring muscles within triceps surae (Laine et al., 2016; Smilde et al., 2016) may play an important role in modulating common synaptic inputs. Consequently, muscle composition and synergistic function likely influence the modulation of common alpha band inputs.

### Common corticospinal input is not affected by changing ankle position

Beta band oscillations in motor unit activity stem from the motor cortex and are mediated by the corticospinal pathway (Brown, 2000; Fisher et al., 2012). The present study found that changes in ankle position did not significantly affect intramuscular nor intermuscular coherence in the beta band. Previous research on corticomuscular coherence showed that beta band oscillations were greater following a large movement than following a small movement (Riddle & Baker, 2006). The minimal modulation of common corticospinal inputs within and across triceps surae muscles observed in the present study may be attributed to the isometric nature of the task, smaller force generated, and focus of the analysis on the stable phase of the contraction.

When pooled across three ankle positions, beta band motor unit coherence within GM was greater than that within other muscles, while SM exhibited the lowest coherence. Since this study did not directly measure activity of the cortex, these differences across muscles need further investigation. A possible explanation is the variation in muscle spindle density and architectural properties within the triceps surae, which may influence sensorimotor integration (Baker, 2007; Witham et al., 2011) (Baker, 2007; Witham et al., 2011), consequently altering cortical oscillations and excitability (Ko et al., 2023).

### Altered neural drive across angles may be explained by inverse patterns of motor unit recruitment threshold for GM and GL

Changes in motor unit recruitment threshold with muscle length have been shown to be modulated by both central command and muscle spindle afferents to the motoneuron (Nishimura & Nakajima, 2002). The present study found motor unit recruitment threshold of GM is greatest compared to the other three muscles at shortened position, and progressively decreased as the ankle shifted from shortened to lengthened position. Similarly, SL exhibited its lowest recruitment threshold in the lengthened position. In contrast, GL exhibited a lower recruitment threshold at shortened position compared to neutral and lengthened positions, potentially as a compensatory strategy to ensure force onset when GM, which contributes significantly to ankle force, is inhibited and not operating on the ascending limb of its force-length curve (Cunnane et al., 2023). Previous studies have also shown greater motor unit recruitment threshold for GM at shortened position compared to lengthened position by altering knee angle while keeping the ankle fixed (Kennedy & Cresswell, 2001; Nishimura & Nakajima, 2002). This may result from reduced inhibitory inputs as the muscle lengthens (Kennedy & Cresswell, 2001; Nishimura & Nakajima, 2002). The opposite trend in GL recruitment threshold may reflect a shift in the activated motoneuron synergy, potentially explaining the increase in motor unit discharge rates for GL and SOL despite changes in coherence found in our previous study (Levine et al., 2023). We also found that recruitment threshold of SM remained lowest and was unaffected by changes in ankle position. This may contribute to the increased neural coupling between GM and SM as muscle length was increased, suggesting a selective central nervous system strategy to enhance shared common inputs between the prime mover and the muscle that is continuously active and less inhibited to maintain force output at lengthened position.

Interestingly, the opposite trend has been observed for the tibialis anterior (TA), with greater recruitment threshold at the lengthened position (Martinez-Valdes et al., 2022; Pasquet et al., 2005). This has been attributed to smaller inhibition of the TA motoneuron pool at shortened positions discussed in the previous section (Pasquet et al., 2005). The change of recruitment threshold observed in the present study is unlikely to be due to shifting of the grid over a lengthened or shortened muscle. The HDsEMG electrode size (96 x 32 mm) was large enough to cover the muscle belly of each triceps surae muscle, allowing us to collect a broad and random sample of motor units from the motor neuron pool. Our findings are also consistent with previous studies that used intramuscular fine-wire electrodes to ensure measurement from the same motor units (Héroux et al., 2014; Nishimura & Nakajima, 2002) and intramuscular needle EMG to record from different positions within the muscle belly (Kennedy & Cresswell, 2001).

## Limitations

To ensure the extraction of high-quality motor unit data from HDsEMG recordings, this study did not involve changes in muscle length under dynamic conditions. Instead, we focused on investigating motor unit coherence during isometric plantarflexions at three different lengths of the triceps surae by altering ankle positions. This conserves the stability of the motor unit action potential shape, which is used to identify motor unit firings. Recent work has expanded HDsEMG decomposition and motor unit editing techniques to slow, controlled dynamic tasks (Glaser & Holobar, 2019; J. Pereira et al., 2024; Oliveira & Negro, 2021; Yokoyama et al., 2021). Future research should explore motor unit coherence during dynamic tasks and daily life activities.

Another limitation is that motor units were not tracked between ankle angles in this study. This can be accomplished by comparing the similarity of motor unit action potential shapes across trials and matching those with a similarity greater than a threshold, as described by (Martinez-Valdes et al., 2017) and was done by (Oliveira & Negro, 2021), for example. Matching motor units resulted in a much smaller number of motor units to be used for subsequent analyses, and as described in the Methods, at least six motor units were required per muscle for intramuscular coherence and at least three per muscle for intermuscular coherence. Therefore, motor units were not matched.

## Conclusion

This study demonstrated how altering ankle position to increase muscle length in an isometric plantarflexion leads to increased shared common inputs across the medial head of the gastrocnemius and the medial compartment of the soleus muscle, particularly in delta and alpha frequency bands. It also provides novel evidence that increased muscle length reduces intramuscular coherence in the alpha band, along with decreases in coefficient of variation of force (i.e., improved force steadiness), and reduced physiological tremor. Additionally, increasing muscle length also results in reduced motor unit recruitment threshold of GM and SL. However, GL exhibited an opposite pattern, showing higher recruitment thresholds at lengthened positions, while SM was not affected by changes in muscle length. This study reveals a neuromuscular control strategy that modulates motor unit recruitment and common synaptic inputs within and shared across the triceps surae to maintain isometric force output at varying muscle lengths. It also highlights the influence of muscle architecture and inhibitory inputs associated with changes in muscle length and the modulation of these inputs by neighboring synergistic muscles. The findings offer new insights into the neural mechanisms underlying muscle coordination and force control in the triceps surae, with potential implications for understanding motor control during multi-joint, dynamic, and more complex tasks.

## Acknowledgements

José L. Pons is supported by the National Science Foundation/National Robotics Initiative under Grant 2024488.

## Author contributions

Experiments were conducted in the laboratory of J.L.P.; J.L.P and J.T.L conceived research; J.L.P and J.T.L designed research. J.T.L performed experiments, X.S.Y analyzed data, interpreted results of experiments, prepared figures, and drafted manuscript; J.T.L assisted results interpretation. X.S.Y, J.T.L and J.L.P edited and revised manuscript. All authors have read and approved the final version of this manuscript.

